# *Rsearch*: An R interface to VSEARCH supporting visualization and parameter tuning

**DOI:** 10.64898/2026.09.10.750626

**Authors:** Cassandra Stamsaas, Torbjørn Rognes, Knut Rudi, Lars Snipen, Hilde Vinje

**Author notes:** Corresponding author: Hilde Vinje. Cassandra Stamsaas, Torbjørn Rognes, Knut Rudi, Lars Snipen.

## Abstract

**Background:** We present *Rsearch*, an R package that integrates the core functionality of VSEARCH into the R environment and extends it with visualization, parameter optimization, and conversion tools for compatibility with other R packages. By making VSEARCH directly accessible in R, *Rsearch* lowers the barrier for using VSEARCH and integrating it with downstream statistical and ecological analyses.

**Results:** Comparative analysis with *DADA2* using mock community data showed that both pipelines produced relative abundance profiles highly correlated with the expected composition. Compared to *DADA2*, *Rsearch* identified fewer OTUs, but these were more consistently prevalent across samples. In contrast, *DADA2* appeared to overestimate diversity by splitting sequences into an excessive number of OTUs. In terms of computational performance, `vs_cluster_unoise` implemented in *Rsearch* was the fastest of all the clustering and denoising methods, while other *Rsearch* functions showed runtimes comparable to *DADA2*. In addition, *Rsearch* provides functions for systematic optimization of trimming and filtering parameters, an important feature for users who may not otherwise have a clear strategy for parameter selection. The package also includes functions to ensure compatibility with other R packages such as *phyloseq*.

**Conclusions:** *Rsearch* offers a practical and accessible framework for analysing metabarcoding data within a single analytical environment and is freely available from The Comprehensive R Archive Network, with the development version hosted on GitHub (https://github.com/CassandraHjo/Rsearch).

## Background

The application of metabarcoding to DNA has revolutionized our ability to study all types of environments, revealing their diversity, structure, and shifts in ecological status [1]. These advances have been made possible by high-throughput sequencing technologies, which enable the rapid sequencing of DNA from diverse environmental samples, together with bioinformatic pipelines [2], [3], [4]. Importantly, metabarcoding is not restricted to microbes but can also be used to investigate fungi, plants, and animals, underscoring its broad applicability across ecosystems and taxa [1].

Several bioinformatics tools have been developed for processing metabarcoding raw data, including QIIME2 [5], MOTHUR [6], *DADA2* [7], Deblur [8], USEARCH [9], VSEARCH [10], and Swarm [11]. These tools differ in terms of underlying algorithms, performance, and user experience, and have been reviewed elsewhere [10], [12], [13]. Some prioritize denoising accuracy, others flexibility, or computational efficiency. VSEARCH stands out as an efficient, multithreaded, open-source tool that supports a complete metabarcoding workflow and is compatible with all major operating systems. Its broad compatibility and speed make it a popular choice in many pipelines, especially when analysing environmental DNA (eDNA), as UNOISE [14] implemented in VSEARCH has demonstrated strong performance in analysing high-diversity environmental samples compared to *DADA2* and Deblur [15]. However, as a command-line-based tool, VSEARCH requires users to be familiar with terminal-based scripting and may be a barrier for many researchers.

To make VSEARCH more user-friendly and accessible to a broader audience, we introduce *Rsearch*, an R-package [16] that integrates VSEARCH into the more user-friendly and widely adopted R environment. Both R and VSEARCH run on all platforms, including UNIX-based, Windows, and macOS. R provides access to a large collection of packages, enabling a wide range of statistical analyses and graphical visualizations. This flexibility allows users to construct flexible workflows tailored to diverse analytical needs. The *Rsearch* package streamlines the full metabarcoding workflow, including trimming and filtering, merging paired-end reads, dereplication, and chimera removal. As R is already widely adopted in metabarcoding research [17], *Rsearch* integration enables researchers to perform end-to-end analysis in a single, cohesive environment. In addition to simplifying access to core VSEARCH functionality, *Rsearch* introduces dedicated functions for data visualization and parameter optimization. These tools provide users with direct support for evaluating sequence quality, selecting trimming and filtering thresholds, and refining analytical workflows based on transparent, reproducible criteria.

### Implementation

This study aimed to make VSEARCH more accessible by developing *Rsearch*, an R package that implements its core functions together with tools for visualization and parameter optimization. The package facilitates integration with common downstream analysis workflows, including *phyloseq* [18]. The *Rsearch* package [19] is freely available at the Comprehensive R Archive Network (CRAN) [20], and the development version can be found on GitHub (https://github.com/CassandraHjo/Rsearch).

The VSEARCH functions implemented in *Rsearch* require that VSEARCH version 2.30.0 or later is installed on the user’s system. Additionally, *Rsearch* includes the *phyloseq* R package [18] as a dependency to facilitate seamless integration with downstream analysis. Installation instructions (including text and video guides) are available on the *Rsearch* GitHub page (https://github.com/CassandraHjo/Rsearch) and on the *Rsearch* website (https://cassandrahjo.github.io/Rsearch/). Once installed, VSEARCH is accessed through R functions, allowing users to utilize VSEARCH’s highly efficient computing capabilities without leaving the R environment.

A core feature of *Rsearch* is the ability to retain VSEARCH output within R’s generic data structures, rather than writing results to external files, as is standard in VSEARCH. This enables users to perform their entire workflow within R, facilitating integration with existing tools for data wrangling and visualization. Alternatively, users may choose to export results as text files, ensuring compatibility with other downstream analysis tools and pipelines that rely on file-based input. Inputs to *Rsearch* functions can similarly be supplied as R data structures or files, allowing flexible integration into both complete workflows and modular use of selected components.

*Rsearch* implements the following VSEARCH commands: cluster_size, cluster_unoise, fastq_join, fastq_mergepairs, fastx_subsample, fastx_filter, fastx_uniques, search_exact, sintax, uchime_denovo, uchime_ref, and usearch_global. Detailed descriptions of these commands are available in the VSEARCH documentation [10], and the mapping between VSEARCH commands and *Rsearch* functions is provided in Supplementary Table S1.

In addition, *Rsearch* introduces new functions that do not exist in VSEARCH. An overview of these functions is presented in this paper, while full documentation, including parameter specifications, is available in the R package documentation and on the website (https://cassandrahjo.github.io/Rsearch/). Functions that wrap or extend VSEARCH functionality, including those for parameter optimization, are prefixed with “vs_” to indicate their dependence on VSEARCH. The new functions are tools for file management (`fastx_combine_files`, `fastx_synchronize`), plotting and visualization (`plot_base_quality`, `plot_read_quality`, `plot_ee_rate_dist`, `plot_size_dist`), read merging optimization (`vs_merging_lengths`, `vs_optimize_truncee_rate`, `vs_optimize_truncqual`), data processing for further analysis (`rsearch_obj`, `rsearch2phyloseq`, `phyloseq2rsearch`), taxonomy (`make_sintax_db`, `taxonomy_distance`, `taxonomy_tree`,`vs_alignment_classification`) and clustering (`vs_cluster_subseq`). Figure 1 provides an overview of a typical *Rsearch* workflow, integrating both VSEARCH-based functions and novel *Rsearch* functionality.

**Figure 1:**
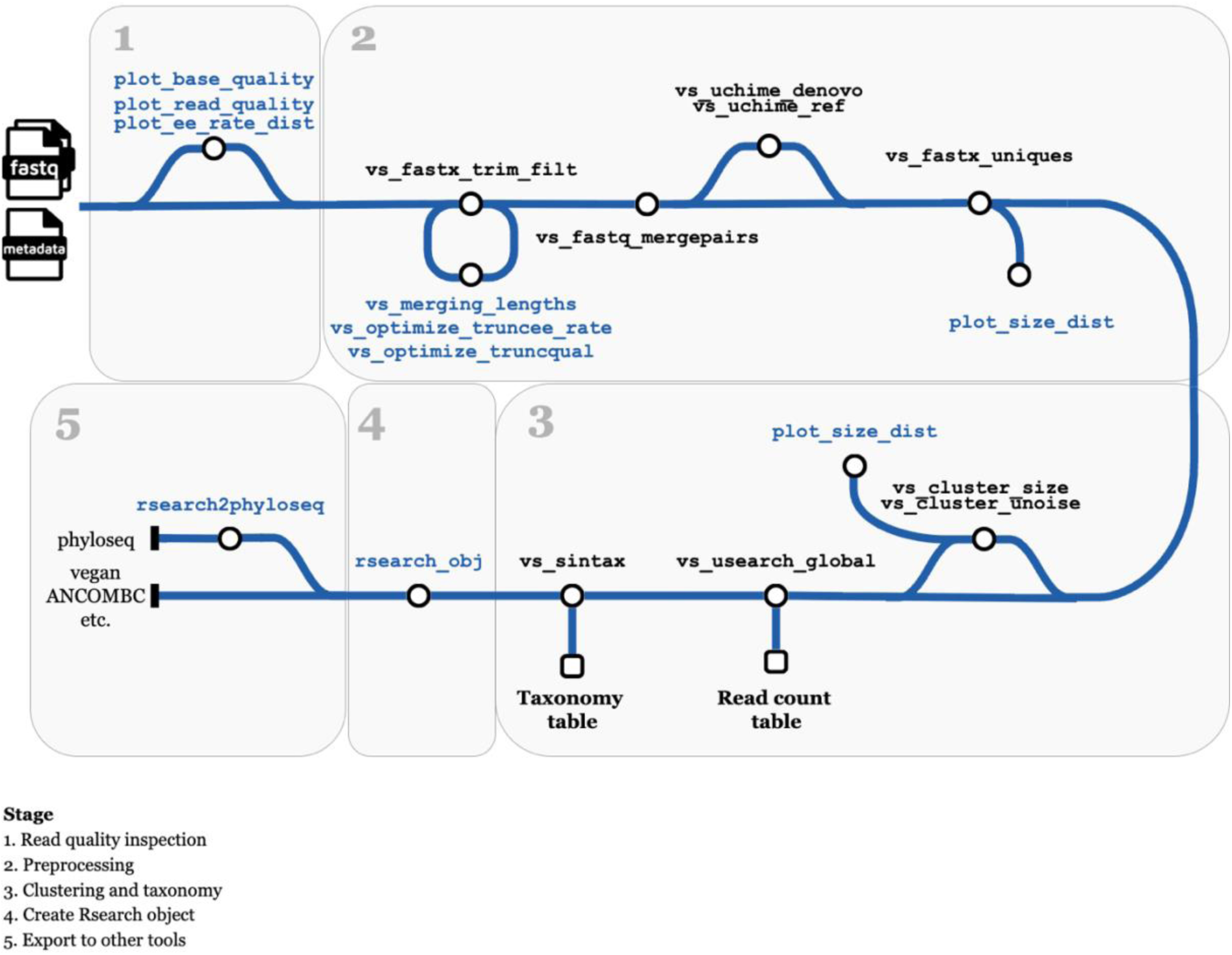
Example workflow for analysing paired-end FASTQ reads with Rsearch. The main analysis path follows a typical metabarcoding pipeline, while the detours represent optional steps that support visualization or optimization of the workflow. The workflow begins with inspecting read quality using dedicated plotting functions. This is followed by trimming and filtering the reads, merging read pairs, removing chimeras, and dereplicating sequences. Trimming can be optimized using two optional functions. The subsequent steps include clustering and taxonomic assignment, after which an Rsearch object is created. This object can be converted into a phyloseq object or exported for use with other tools and packages. The new functions included in Rsearch are shown in blue, while functions using existing VSEARCH functionality are shown in black and always starts with “vs_”.

### New functions provided by *Rsearch*

*Rsearch* provides two functions for file management in metabarcoding workflows. `fastx_combine_files` combines all sequence files of a given type in a directory into one file or table, facilitating aggregation before clustering or taxonomic classification. `fastx_synchronize` synchronizes paired FASTA/FASTQ files or tables, ensuring that the forward (R1) and reverse (R2) reads remain correctly matched in case these files have been processed separately, resulting in an unequal number of reads.

*Rsearch* provides several plotting functions to support data exploration. These visualize key quality and distribution metrics from sequencing data. `plot_base_quality` displays mean and median quality scores at each base position with error bars and optionally adds a shaded box representing the expected overlap length from read merging (Figure 2). This merging step is computed internally, adding minimal runtime while providing useful insight into overlap quality. `plot_read_quality` summarizes read quality by plotting either expected error (EE) rate or average quality score against read length.

**Figure 2:**
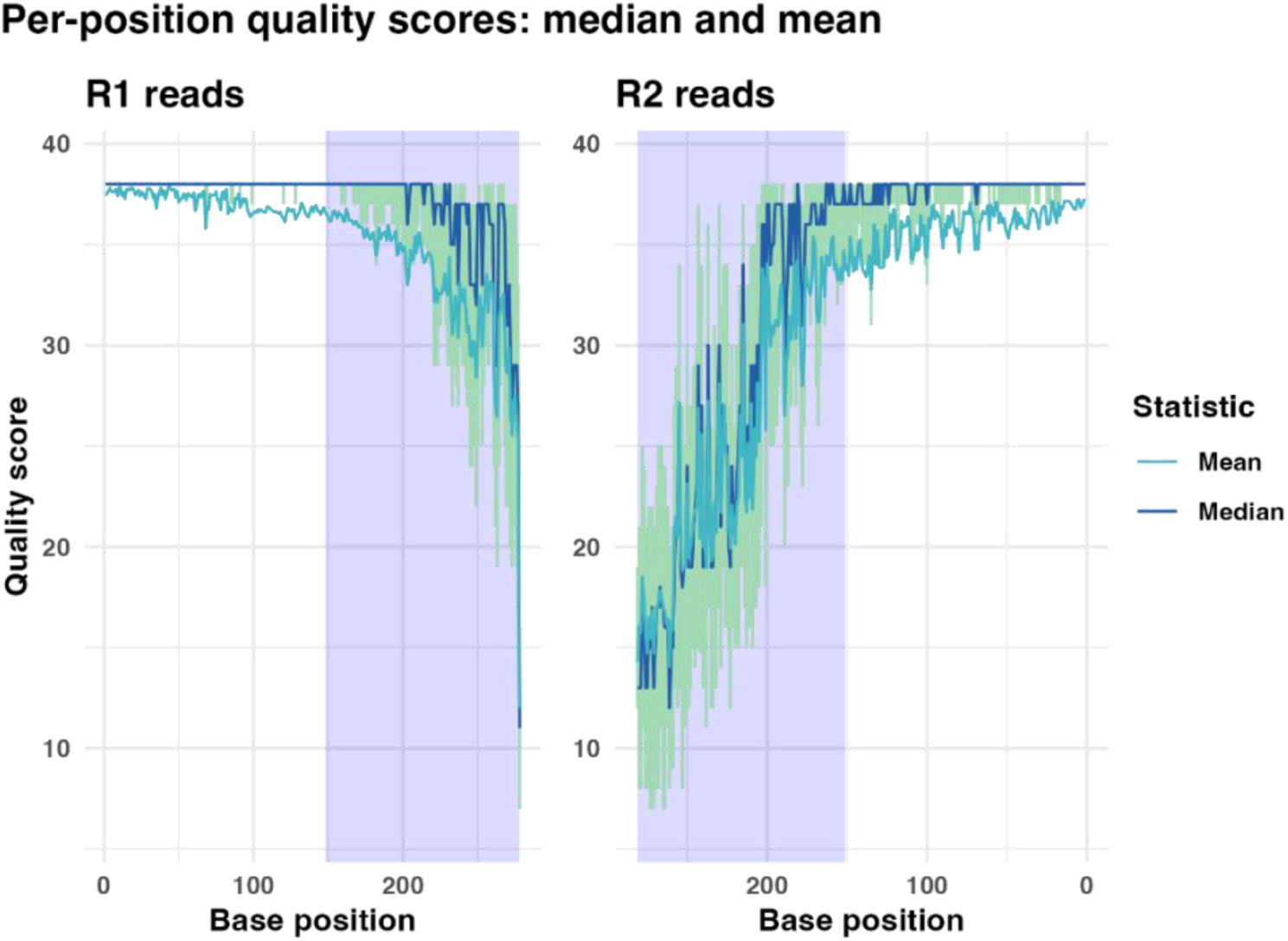
`plot_base_quality` returns a ggplot object showing base quality scores by position across reads. The x-axis indicates the base position within the reads, while the y-axis shows the corresponding quality scores. Light blue line indicates the mean quality score for the given position and the dark blue line indicates the median. The shaded area indicates the average overlap length of the read pairs after merging.

`plot_ee_rate_dist` shows the distribution of EE rates across reads, and `plot_size_dist` visualizes read abundance distributions. All plotting functions return *ggplot* objects, allowing users to further customize the visualizations as needed. This design choice leverages the flexibility and familiarity of the *ggplot2* system, making it easier to tailor plots to specific analytical or presentation needs - an important strength for users who wish to integrate *Rsearch* outputs into broader R-based workflows.

Merging paired-end reads is a critical preprocessing step, but selecting appropriate trimming and filtering parameters can be challenging. *Rsearch* provides three functions to evaluate and optimize merging outcomes. `vs_merging_lengths` reports statistics on forward, reverse, merged, and overlap lengths, and generates a plot of these metrics (Figure 3). `vs_optimize_truncee_rate` identifies optimal thresholds for the `truncee_rate` parameter, which truncates the 3’ end of reads so that their average expected error per base does not exceed a specified value. The function tests a user-defined range and selects the setting that maximizes merged read pairs above a copy number threshold (default 2). The optimization is based on the assumption that sequencing errors are more likely to occur in singletons, therefore the focus is on merged reads with a copy number of 2 or more. A higher number of these high-quality merged sequences is taken as an indicator of better merging performance. Reads with average expected error greater than 0.01 (default) are filtered out.

**Figure 3:**
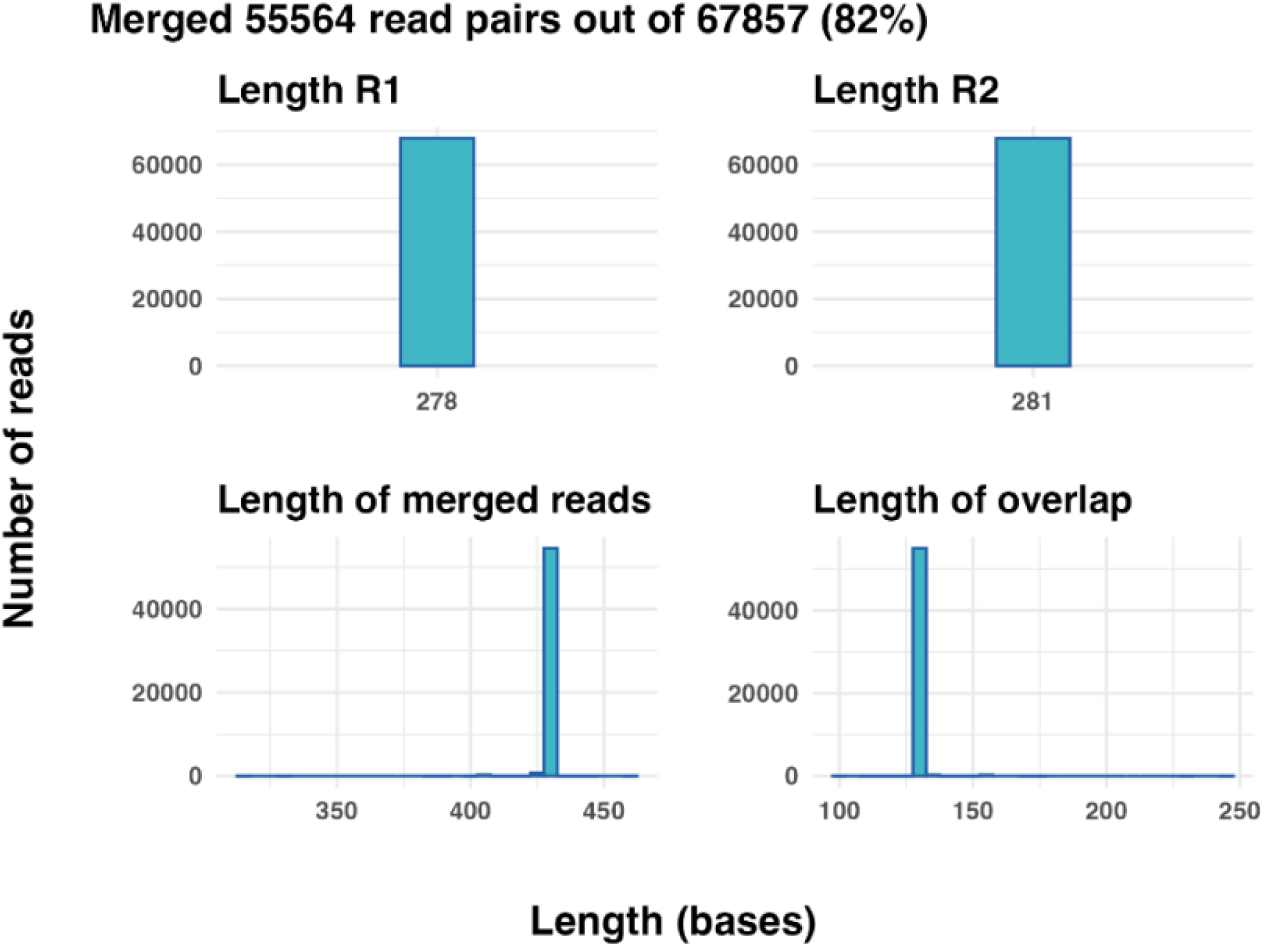
`vs_merging_lengths` returns a ggplot object with four panels visualizing different length statistics during merging. The x-axis shows the length in bases, and the y-axis shows the number of reads with the given length. The plot title shows the number of read pairs that were merged.

This value corresponds to a Phred quality score of 20, a commonly used threshold for defining high-quality reads. The function outputs a summary plot (Figure 3A): the top panel shows merged read counts above the threshold at each `truncee_rate` value, with a red dashed line marking the optimal value, and the bottom panel shows average forward and reverse read lengths. `vs_optimize_truncqual` applies the same optimization procedure to the `truncqual` parameter, which trims reads so that their total expected error does not exceed a specified value (Figure 3B). This supports an informed selection of `truncee_rate` or `truncqual` to improve merging performance.

To support downstream analysis, *Rsearch* provides data processing functions centred around the `rsearch_obj` function. This function organizes read counts, sequences and sample data into a structured *Rsearch* object.

Given the widespread use of *phyloseq* [18] in microbiome studies (see Wen et al. [17] for a broader review), *Rsearch* ensures compatibility through conversion functions. `rsearch2phyloseq` transforms an *Rsearch* object into a *phyloseq* object, enabling easy access to *phyloseq* tools after preprocessing. Conversely, `phyloseq2rsearch` converts back to an *Rsearch* object, allowing for seamless exchange between the two frameworks. For taxonomy-focused analyses, *Rsearch* provides two functions. `make_sintax_db` generates a Sintax-formated FASTA file from a user-supplied data frame of headers, sequences, and taxonomic classifications, enabling the creation of custom reference databases without relying on existing ones. `taxonomy_tree` builds a phylogenetic tree by inferring Operational Taxonomic Unit (OTU) relatedness from shared taxonomic ranks, using a distance metric derived from taxonomic depth.

### A worked example and comparison with *DADA2* using mock community data

To demonstrate the use of *Rsearch* we process a mock community dataset with 16S rRNA gene data. To show that it performs well, we have carried out the same analysis using the well-established R package *DADA2* [7]. A mock dataset has the advantage that we already know the correct outcome in advance. That is, we know exactly which taxa are present and at which relative abundance, and it therefore allows us to more reliably evaluate the results we obtain.

The mock community dataset used in this study was generated by amplifying the ZymoBIOMICS Microbial Community Standard (Zymo Research Corporation, California, USA. Product D6300) with primers targeting the V3-V4 region of the 16S rRNA gene. Sample handling, DNA extraction, and sequencing protocols are described in detail in Philip et al. [21]. The mock community contained eight bacterial species represented by 10 distinct 16S rRNA gene sequences since two bacteria contained two 16S rRNA gene variants in their genomes. Details of the initial processing steps, including demultiplexing, barcode removal, and primer trimming, are described in detail in Nilsen et al. [15].

For consistency, we refer to the resulting representative sequences from both pipelines as OTUs, regardless of how they were obtained. For both pipelines, trimming thresholds were systematically evaluated to optimize merging efficiency. In *Rsearch*, the 3’ end trimming was optimized using the `vs_optimize_truncee_rate` function with default settings. The optimal value was then applied via `vs_fastx_trim_filt`, along with `minlen` = 20 and `maxee_rate` = 0.01.

In *DADA2*, 3’ end trimming was optimized manually (there is no built-in tool for it) by testing different values of `truncLen` in the `filterAndTrim` function. Several combinations of trimming thresholds for R1 and R2 reads were evaluated to modulate the overlap length, which affects merging success. Trimming lengths were varied to generate approximate overlaps of 60 base pairs (bp), 40 bp, 20 bp, 12 bp (the default minimum in *DADA2*), and below 12 bp, to assess the impact on merging outcomes.

In *Rsearch*, the analysis proceeded with merging (`vs_fastq_mergepairs`), dereplication (`vs_fastx_uniques`), and chimera removal (`vs_uchime_denovo`). All reads were pooled into a single file and dereplicated wit `minsize` = 2 to remove singleton reads before clustering. OTU clustering was then performed using two approaches: `vs_cluster_size` across multiple identity thresholds (90-99%), `vs_cluster_unoise` with default parameters. For `vs_cluster_size`, the same identity threshold was applied in `vs_usearch_global` to generate the read count tables, whereas for `vs_cluster_unoise`. `vs_usearch_global` was run at 97% identity to produce the read count table.

In *DADA2*, trimmed reads were used to learn error rates (learnErrors), followed by dereplication, denoising (dada), read merging (mergePairs), and chimera removal (removeBimeraDenovo).

Since both pipelines were run on identical input data, their outputs could be directly compared in terms of microbial composition, number of OTUs and reads, OTU prevalence, over-merging, and over-splitting. Microbial composition was assessed by aligning the centroid sequence of each OTU to the reference sequences of the mock community using the `vs_usearch_global` function with a 99% identity threshold, retaining only the best matching OTU for each reference sequence. Read counts were aggregated per OTU, and relative abundances were calculated as the proportion of reads per OTU relative to the total number of reads. The resulting profiles were compared to the expected relative abundances of the mock community using Pearson correlation coefficients. In addition to taxonomic accuracy, we evaluated the total number of OTUs detected by each method and analysed their prevalence across the 15 samples. Read retention was tracked throughout the pipelines to quantify the proportion of sequences retained after filtering, merging, chimera removal, and other processing steps.

To assess over-merging and over-splitting, OTU centroids were mapped to the reference sequences using the `vs_usearch_global` function with a 99% identity threshold, retaining all matches rather than only the best hit. Over-merging was defined as multiple reference sequences being grouped into a single OTU (i.e., some reference sequences were not recovered). Over-splitting was defined as a single reference sequence being split across multiple OTUs.

### Runtime analysis

Runtime was measured using the `Sys.time` function in R [16]. The new *Rsearch* functions were evaluated over five iterations, each using a FASTQ dataset of approximately 68 000 paired-end reads. For analysis of complete pipelines, the mock community dataset comprising approximately 1.4 million read pairs, with a median of 75 000 read pairs per sample, was used. To provide a fair comparison, pipelines were executed with a single thread, as multithreading is not supported by *DADA2* on Windows systems. All benchmarks were performed in R version 4.5.1 on an Apple Silicon (ARM64) platform running macOS Sequoia 15.6.1 and Windows 11 Enterprise system with a 13th Gen Intel® Core™ i5-1345U processor (1.60 GHz), 16 GB RAM, 64-bit operating system.The pipelines included typical steps in metabarcoding analysis, such as filtering and trimming, merging, dereplication, chimera removal, and clustering or denoising, and produced a final read count table as output.

## Results

The main outcome of this work is the development of the open access R package *Rsearch*, which provides new functionality for OTU analysis. To demonstrate the new functions implemented in *Rsearch*, we present example outputs generated using the mock community data. These examples highlight the added functionality and flexibility of the package. Detailed guidance on integrating these functions into complete workflows is provided in the package tutorial (https://cassandrahjo.github.io/Rsearch/articles/tutorial.html).

### New functions provided by *Rsearch*

The `plot_base_quality` function is used to assess the quality of FASTQ reads and returns a *ggplot* object (Figure 2), allowing for further customization. The plot displays mean (turquoise line) and median (dark blue line) base quality scores across all positions in the reads. Base positions are shown on the x-axis, and quality scores on the y-axis. Green error bars represent the interquartile range (25th to 75th percentiles). To reflect the fact that R1 and R2 reads are sequenced in opposite directions, the x-axis for the R2 panel is flipped, making the plot easier to interpret. Blue shaded regions indicate the average overlap length resulting from merging the paired-end reads, providing insight into base quality within these overlapping segments. All optional parameters in this function have default values, allowing the plot to be generated by specifying only the input data. However, several parameters enable customization: error bars can be adjusted to different quantile ranges, the plot title can be modified or removed, quality statistics can be toggled on or off, and the shaded overlap box can be activated. The resulting plot offers a clear visual summary of base quality across read positions, helping to guide decisions about whether and how to trim the reads. It can also be used after trimming and filtering to assess changes in quality and to evaluate the quality within the overlap region.

During runtime analysis, `plot_base_quality` demonstrated consistent performance. When the shaded overlap box was activated, the median runtime was 2.2 seconds. Without the shaded box, the median runtime was 1.8 seconds.

The `vs_merging_lengths` function is used to compute length statistics of paired-end reads and the resulting merged reads and returns a *ggplot* object (Figure 3). The plot displays the length of the R1, R2 and merged reads and the length of the overlap across all reads. Length in number of bases is shown on the x-axis, and number of reads on the y-axis. All optional parameters in this function have default values, allowing the plot to be generated by specifying only the input data.

During runtime analysis, vs_merging_lengths demonstrated consistent performance, with a median runtime of 2.8 seconds.

The functions `vs_optimize_truncqual` and `vs_optimize_truncee_rate` are used to identify optimal trimming thresholds for maximizing merging performance. Each function returns both a results table and a *ggplot* object visualizing the results (Figure 4). The top panel of each subplot displays the number of merged read pairs at each tested `truncqual` or `truncee_rate` value (light green line), while the bottom panel shows the average read lengths of R1 and R2 reads (turquoise and dark blue lines, respectively). The x-axis represents the range of trimming parameter values tested.

**Figure 4:**
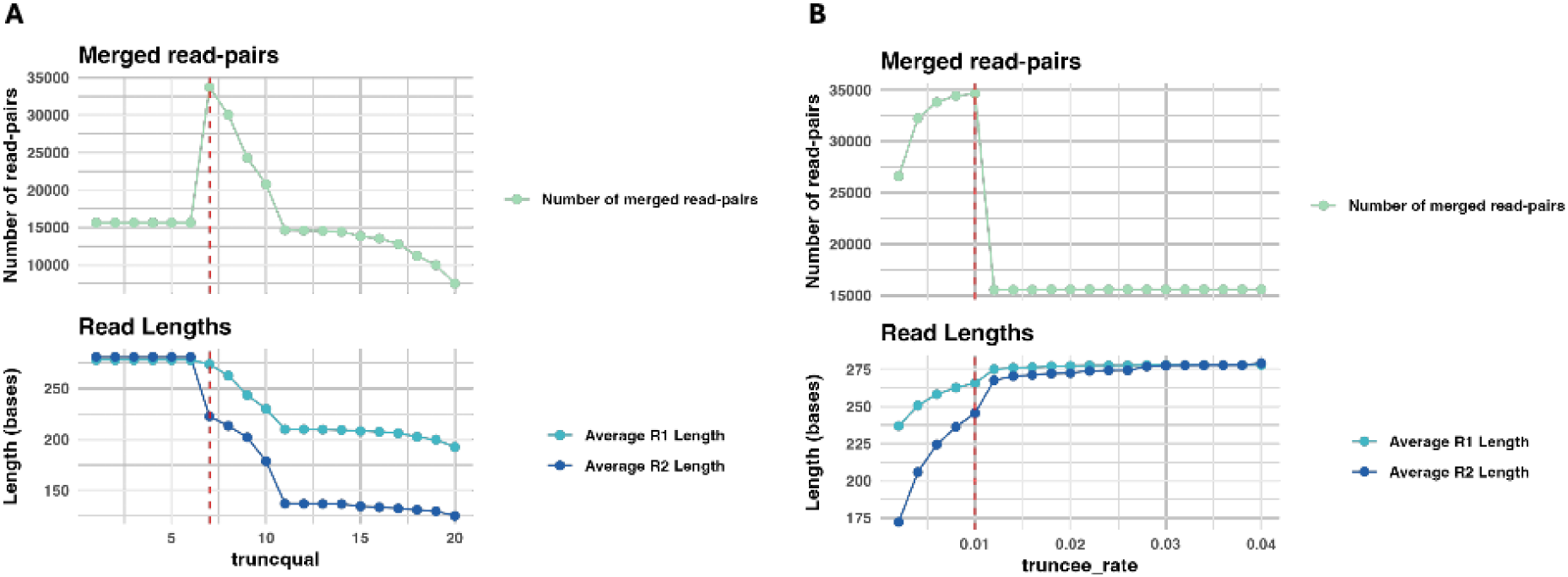
Merging performance across quality filtering parameters. In each plot, the top panel shows the number of successfully merged read pairs (light green line) with a copy number greater than 2 and a minimum overlap of 10 bases for each parameter value. The bottom panel shows the average read lengths of R1 (turquoise line) and R2 (dark blue line). The x-axis indicates the tested range of either `truncqual` or `truncee_rate` values, and the red vertical line marks the optimal value identified by the function. (**A**) `vs_optimize_truncqual` results across `truncqual` values from 1 to 20, with 7 identified as optimal. (**B**) `vs_optimize_truncee_rate` results across `truncee_rate` values from 0.002 to 0.04, with 0.01 identified as optimal.

All optional parameters have default values, allowing the function to run with only the input data specified. However, adjusting the range of the trimming parameter may improve the likelihood of identifying the optimal setting.

Both functions demonstrated consistent performance during runtime analysis. `vs_optimize_truncqual` had a median runtime of 47.8 seconds and `vs_optimize_truncee_rate` had a median runtime of 49.8 seconds.

### A worked example and comparison with *DADA2* using mock community data

Trimming thresholds were evaluated with `vs_optimize_truncee_rate` to identify settings that maximize read merging. A baseline from raw reads using `vs_merging_lengths` showed that more than 80% of read pairs were successfully merged across all samples, with some approaching 90%. Applying `vs_optimize_truncee_rate` to the first sample from each sequencing run consistently identifying 0.01 as the optimal `truncee_rate`. Results for one sequencing run are shown in Supplementary Figure S1.

Using this threshold, reads were trimmed, merged, dereplicated, and filtered for chimeras. All reads were then pooled across samples, dereplicated again, and clustered with `vs_cluster_size` at 90 to 99% identity and with `vs_cluster_unoise`, generating OTUs and read count tables. Table 1 summarizes read retention after each step in the pipeline. Approximately 1 100 000 reads remained after processing with *Rsearch*, with about 300 000 reads discarded in total, the largest fraction removed during chimera removal.

**Table 1:** Summary of read tracking information for Rsearch and DADA2 applied to the mock community data.

| Pipeline | Total no. of input reads | Total no. of reads after: |  |  |
| --- | --- | --- | --- | --- |
|  |  | Trimming | Merging | Chimera removal |
| ***Rsearch*** | 1 382 865 | 1 289 083 | 1 222 140 | 1 080 631 |
| ***DADA2*** | 1 382 865 | 1 380 859 | 1 312 917 | 1 127 958 |

To mirror the optimization approach available in *Rsearch*, we systematically evaluated the impact of trimming length on the merging performance of *DADA2*. Using initial `truncLen` values of 250 (R1) and 220 (R2), the average overlap was approximately 40 bp. Additional parameter settings extended the overlap to about 60 bp (260/230) or reduced it to below the *DADA2* default threshold of 12 bp (230/200). Merging success was largely unaffected as long as the overlap exceeded 12 bp (Supplementary Figure S2), demonstrating that a `truncLen` setting of 250/220 was sufficient to maximize merged read pairs in this dataset. The corresponding read tracking information are presented in Table 1. Following this pipeline, approximately 1 100 000 reads were retained, while about 300 000 reads were discarded, with the largest fraction removed during chimera removal.

OTU generation was also evaluated across trimming thresholds in *DADA2*. When overlap exceeded 12 bp, OTU counts before chimera removal ranged from about 3 400 to 4 100, but chimera removal consistently reduced this to roughly 3% of the initial number (Supplementary Table S2). With `truncLen` set to 250/220, *DADA2* produced 3 977 initial OTUs, of which 121 remained after chimera removal (Table 3).

**Table 2:** Pearson correlation between observed and expected relative abundances in the mock community data.

| Pipeline | Clustering/denoising function | Identity threshold | Correlation |
| --- | --- | --- | --- |
| *Rsearch* | `vs_cluster_size` | 90% | 0.6971 |
|  |  | 95% | 0.9932 |
|  |  | 96% | 0.9932 |
|  |  | 97% | 0.9933 |
|  |  | 98% | 0.9933 |
|  |  | 99% | 0.9944 |
|  | `vs_cluster_unoise` |  | 0.9878 |
| *DADA2* | `dada` |  | 0.9937 |

Correlations are shown for each clustering or denoising approach. Values in the clustering function column indicate the applied identity threshold.

**Table 3:**
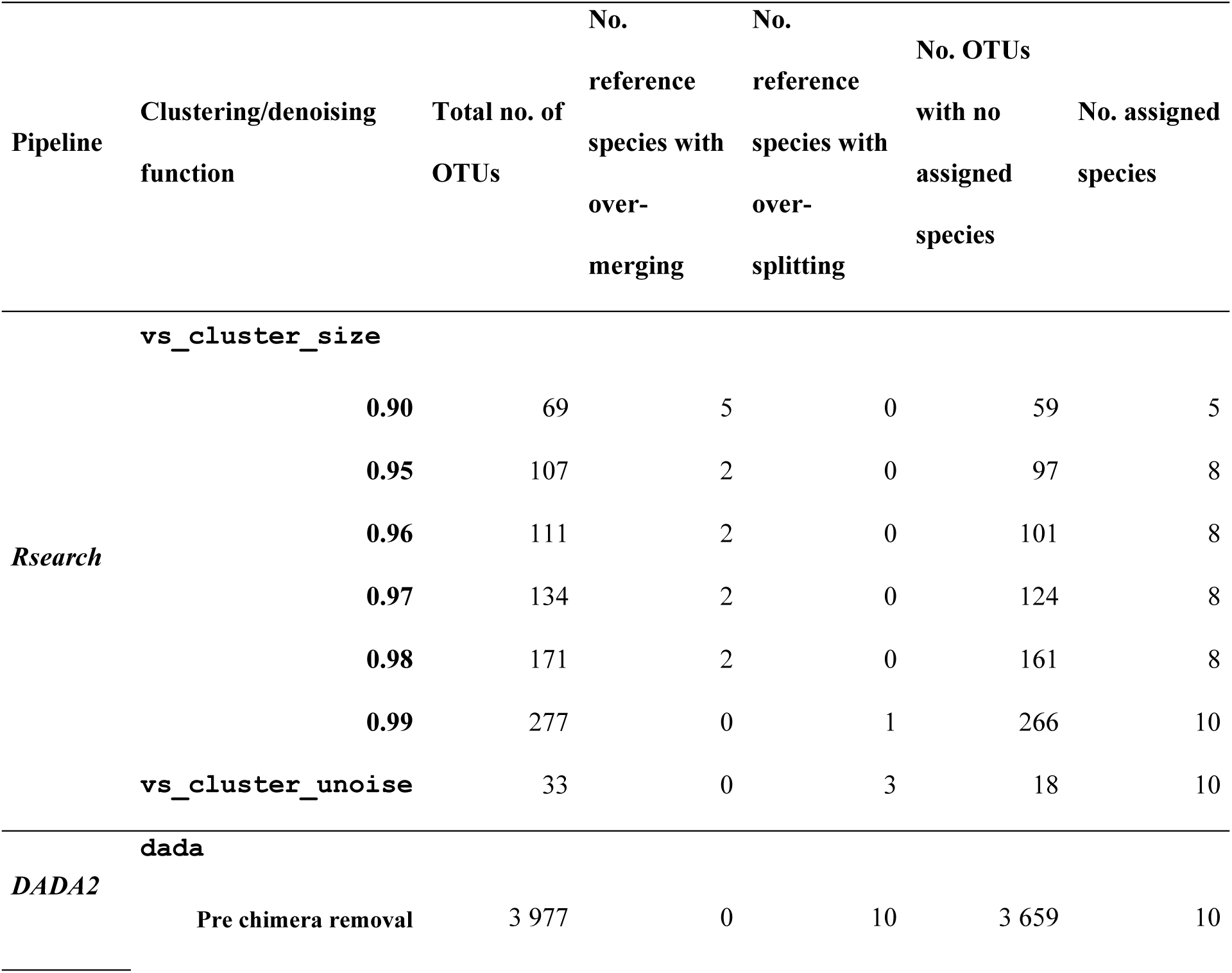
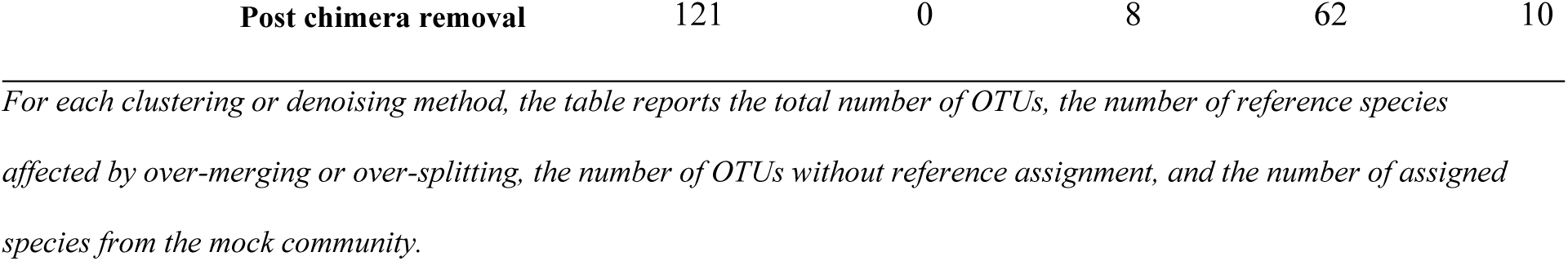
Summary of over-merging and over-splitting in Rsearch and DADA2 on mock community data.

To assess accuracy, results from *Rsearch* and *DADA2* were compared to the known composition of the mock community. Centroid sequences from each clustering or denoising approach were aligned to the reference sequences, and similarity was evaluated by correlating the observed and expected relative abundance profiles. For *Rsearch*, all `vs_cluster_size` identity thresholds above 90%, as well as `vs_cluster_unoise`, showed strong agreement with the expected composition, with correlation coefficients near 0.99. *DADA2* achieved the same level of agreement, indicating comparable performance (Table 2).

Figure 5 presents observed versus expected relative abundances for *Rsearch* at 99% clustering identity using `vs_cluster_size` and for *DADA2* after chimera removal. The relationships were approximately linear, with only minimal differences between the two pipelines.

**Figure 5:**
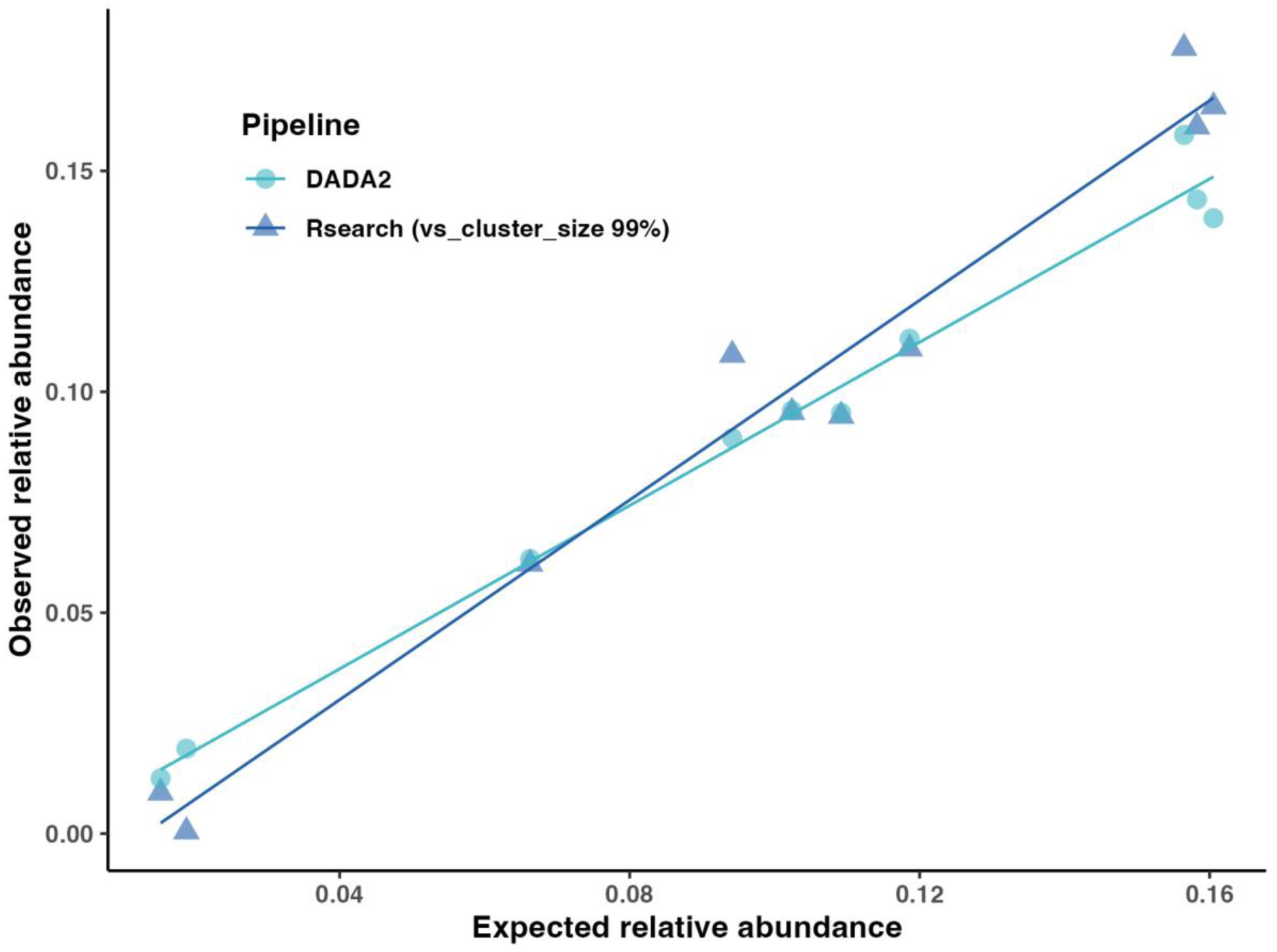
Observed versus expected relative abundances in the mock community. Scatterplot of expected relative abundances (x-axis) versus observed relative abundances (y-axis) inferred by Rsearch at 99% identity in `vs_cluster_size(`dark blue triangles) and by DADA2 (light blue circles). Each point represents one taxon. Lines indicate least-squares linear fits.

The number and prevalence of OTUs were also examined. The mock community data set contained the same eight taxa (10 16S rRNA gene variants) in all 15 samples, so any prevalence below 15 indicates an artefactual OTU. Figure 6 shows the distribution of OTU prevalence across samples, with the horizontal dashed line marking the ideal outcome, where ten OTUs are detected in all 15 samples and no OTUs are detected in fewer samples. OTU prevalence, defined as the number of samples in which an OTU was detected, is shown in Figure 6. In *Rsearch*, OTU counts increased with higher identity thresholds in `vs_cluster_size`, reflecting more stringent clustering, while `vs_cluster_unoise` produced 33 OTUs (Table 3). All clustering methods generated more OTUs with prevalence 15 than with prevalence 1, while `vs_cluster_unoise` produced about the same number of OTUs with prevalence 1 and 15.

**Figure 6:**
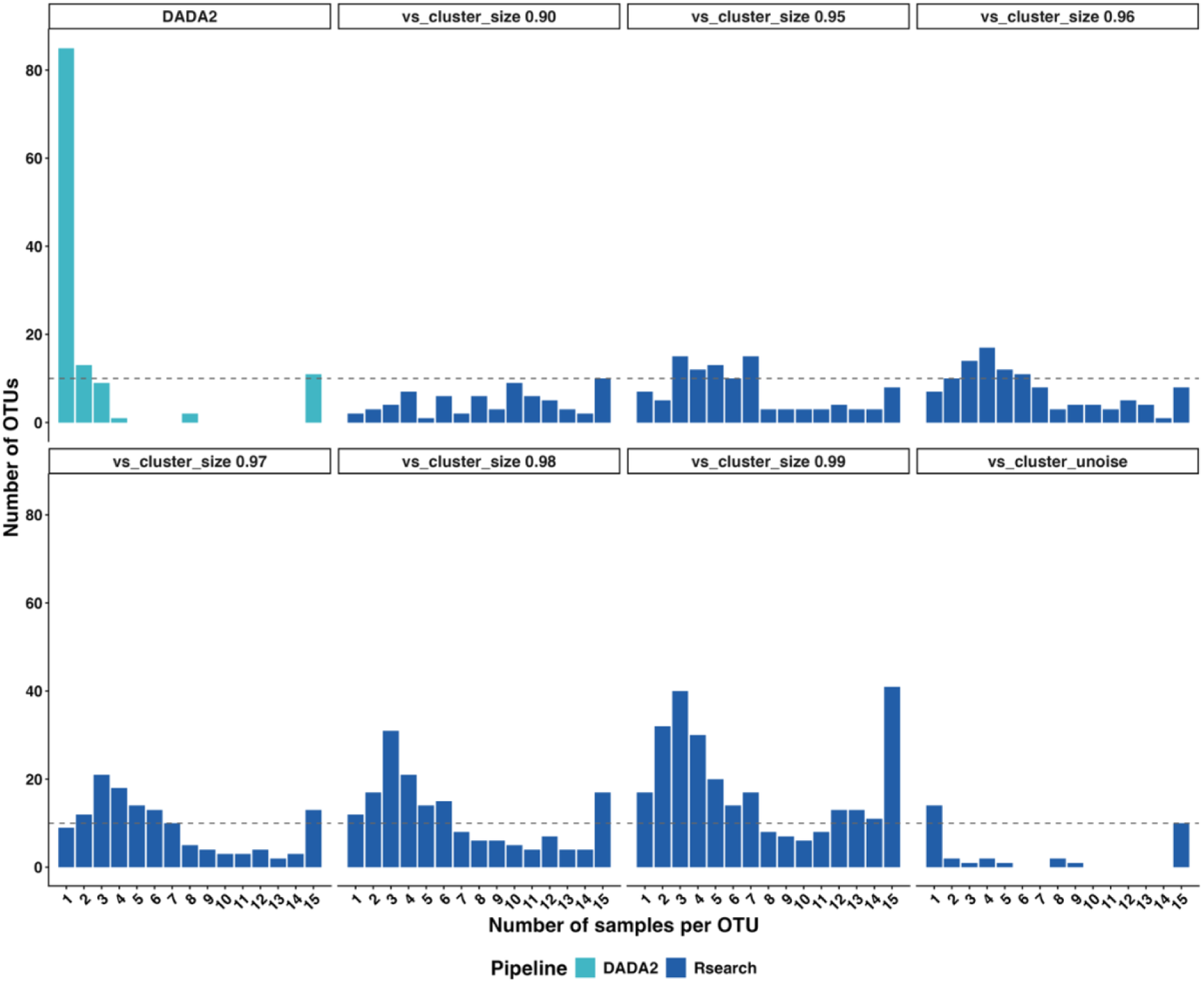
Distribution of OTU sample prevalence across all samples. The x-axis indicates the number of samples in which each OTU was detected (non-zero read count), and the y-axis shows the number of OTUs with that prevalence. Dark blue bars represent Rsearch results, and light blue bars represent DADA2 results. The horizontal dashed line indicates the expected number of OTUs (10) present in all 15 samples.

In *DADA2,* OTU prevalence was more variable, with most OTUs detected in only one sample and 11 OTUs detected across all 15 samples (Figure 6). Given the known composition of the mock community, all clustering and denoising methods appeared sufficient to capture the expected diversity, as each consistently produced at least eight OTUs present across all samples.

Table 3 summarizes the extent of over-merging and over-splitting across clustering and denoising methods. Over-merging was defined as collapsing multiple reference sequences into a single OTU, and over-splitting as dividing a single reference sequence into multiple OTUs. *Rsearch* showed consistently low over-splitting, with no cases across `vs_cluster_size` thresholds except at 99% identity, where one instance occurred; `vs_cluster_unoise` over-split three. Over-merging in `vs_cluster_size` decreased with increasing identity and stabilized between 95 and 98%, while `vs_cluster_unoise` showed no over-merging. *DADA2* showed no over-merging but over-split all 10 reference sequences prior to chimera removal and eight after. All pipelines generated a substantial fraction of unassigned OTUs.

Computational performance was assessed by running all pipelines on the 15 mock community samples. Across five runs, `vs_cluster_unoise` showed the shortest runtimes. `vs_cluster_size` exhibited runtimes comparable to *DADA2*, with longer runtimes at higher identity thresholds. *DADA2* performance varied by platform, with shorter runtimes on macOS than on Windows.

## Discussion

In this work, we introduce *Rsearch*, an R package that implements the core functionality of VSEARCH within the R environment. By providing VSEARCH’s methods directly in R, the package makes these tools more accessible and user-friendly, while also lowering the barrier for integration with downstream statistical and ecological analyses.

In addition to implementing existing VSEARCH commands, *Rsearch* introduces new functions that support different phases of metagenome analysis. Among these are plotting functions, such as `plot_base_quality`, which facilitate quality assessment of sequencing reads. This function visualizes per-base read quality with summary statistics and error bars, enabling informed decisions about trimming strategies. When applied to paired-end data, the function produces dual panels with shaded overlap regions, illustrating how read pairs would merge under current settings. Another relevant function, `vs_merging_lengths`, reports and plots statistics on read and overlap lengths during the merging process.

In addition to visualization, *Rsearch* provides tools for parameter optimization. Functions such as `vs_optimize_truncqual` and `vs_optimize_truncee_rate` help users identify effective trimming and filtering parameters. Parameter choice is often challenging, and many users default to preset values. These optimization functions allow systematic testing of thresholds and provide outputs in both customizable plots and summary tables, thereby supporting transparent and reproducible parameter selection.

A natural comparison for *Rsearch* is *DADA2*, another widely used R-based tool for metabarcoding data analysis. Although both pipelines aim to infer biological sequence variants, they differ in key processing steps, particularly in how and when read merging is performed. *Rsearch* merges read pairs early, thereby preserving pairwise information, whereas *DADA2* merges only after denoising R1 and R2 reads separately. To validate that *Rsearch* performs reliably and generates results consistent with established methods, we compared it with *DADA2* using mock community data, as has also been done in previous evaluations of VSEARCH and *DADA2* (i.e. [22], [23]). This approach enabled direct assessment of read retention, OTU recovery, computational efficiency, and agreement with the known community composition.

The comparison revealed both similarities and differences. Both pipelines produced relative abundance profiles that were highly correlated with the expected composition, indicating accurate taxonomic representation. *DADA2* initially generated a substantially larger number of OTUs, consistent with previous studies [24]. Most of these OTUs were subsequently removed during chimera removal, and many of the retained OTUs showed low prevalence, often occurring in only a single sample. This outcome was unexpected in the context of a mock community, where identical composition was present in all samples. We found no explanation for the large number of OTUs removed in the available *DADA2* documentation or source code. In contrast, *Rsearch* generally produced fewer and more prevalent OTUs, while still recovering the majority of expected species.

All methods, in both *Rsearch* and *DADA2*, generated a substantial fraction of unassigned OTUs when assessing over-splitting and over-merging. In this study, only the reference sequences of the mock community were used as the reference database for taxonomic assignment. Assignment was performed with `vs_usearch_global` at a 99% identity threshold. The high proportion of unassigned OTUs may reflect contamination in the samples or sequencing errors that produced centroid sequences too different from the reference sequences to be matched during alignment.

Differences also emerged when examining over-merging and over-splitting. *DADA2* showed no over-merging but consistently over-split 8 to 10 of the 10 reference species. *Rsearch*, by contrast, exhibited minimal over-splitting across clustering and denoising methods, with some consistent over-merging. Consequently, *DADA2* tends to inflate diversity, whereas *Rsearch* is more conservative.

Performance metrics further distinguished the two pipelines. `vs_cluster_unoise` in *Rsearch* was the fastest pipeline, while `vs_cluster_size` and *DADA2* showed comparable runtimes. Direct execution of VSEARCH in the terminal revealed an overhead from the R implementation. Implementing VSEARCH into R improves accessibility and enables visualization and downstream analyses within a single environment, but at the cost of runtime efficiency.

Another important distinction between the two pipelines is the range of analytical strategies they support. In *Rsearch*, users can choose between denoising and identity-based clustering, whereas *DADA2* only implements denoising. This flexibility is advantageous, as the ever-improving sequencing technologies means the technical noise will eventually be ignorable. But even with diminishing technical noise, biological noise will remain and will always require some degree of clustering [25].

Sampling eDNA often targets microbial communities with high diversity, and this places particularly high demands on data processing. Datasets are typically large, complex, and heterogeneous, making reliable and transparent pipelines essential to ensure reproducibility and comparability across studies. An important consideration in this context is the balance between over-splitting and over-merging. In high-diversity environments, over-splitting can lead to severely inflated diversity estimates with many low-prevalent OTUs detected in only a few samples, which complicates ecological interpretation. For applications such as tracking community changes across time or space, it is preferable to obtain fewer but more stable OTUs. In this context, denoising approaches such as UNOISE implemented in VSEARCH (and *Rsearch*) have been shown to perform well in high diversity datasets [15]. By integrating VSEARCH into R and extending it with visualization, optimization, and conversion functions, *Rsearch* provides a practical framework for processing metabarcoding data within a single analytical environment. Continued development and benchmarking of such tools will be critical to ensure robust handling of eDNA metabarcoding data.

## Conclusions

We have implemented *Rsearch*, an R package that integrates the core functionality of VSEARCH into the R environment and extends it with tools for visualization, parameter optimization, and easy handover to other R packages. Comparison with *DADA2* showed that *Rsearch* produces results consistent with established methods while offering complementary strengths, including the option to apply either denoising or clustering, or both, and efficient runtimes for selected methods. These features make *Rsearch* a practical and accessible framework for metabarcoding analyses within a single analytical environment.

## Declarations

### Ethics approval and consent to participate

Not applicable.

## Consent for publication

Not applicable.

## Availability of data and material

The R package is freely available from The Comprehensive R Archive Network [20]. It is most easily obtained by starting R and running `install.packages(“Rsearch”)`in the console window. The development version is available on GitHub (https://github.com/CassandraHjo/Rsearch). Descriptions on how to download the development version of the package is provided in the README file on GitHub. The dataset (raw sequencing data) supporting the conclusions of this article is available in the NCBI Sequence Read Archive (SRA) under the accession number PRJNA1321533. All scripts and related files are available at https://github.com/CassandraHjo/Rsearch-supplementary-scripts.

## Competing interests

The authors declare that they have no competing interests.

## Funding

This paper is a part of the PhD-project of CS, and this project has been 100% financed by the Norwegian University of Life Sciences.

## Authors’ contributions

Authors CS, TR, LS and HV have all contributed significantly to the programming and documentation of the software. All authors have contributed to the preparation and writing of this manuscript. All authors have read and approved the final manuscript.

## Acknowledgements

We would like to thank Senior Engineers Tonje Nilsen and Inga Leena Angell for their technical assistance with the laboratory analyses and data processing.

## Availability and Requirements

**Project name:** Rsearch

**Project home page:** https://cassandrahjo.github.io/Rsearch/

**Operating system(s):** Platform independent

**Programming language:** R

**Other requirements:** VSEARCH version 2.30.0 or later, and the phyloseq R package

**License:** GNU GPL

**Any restrictions to use by non-academics:** None

## Additional files

File name: Supplementary material

File format: .docx

Title: Supplementary material

Description: Supplementary material containing both figures and tables.

## Notes

https://github.com/CassandraHjo/Rsearch

https://github.com/CassandraHjo/Rsearch-supplementary-scripts

### Competing Interest Statement

The authors have declared no competing interest.

