## Supplementary material for "*Rsearch*: An R interface to VSEARCH supporting visualization and parameter tuning"

### Supplementary materials

#### Supplementary Tables

*Table S1: Mapping between VSEARCH commands and Rsearch functions.*

| VSEARCH command | *Rsearch* function |
|---|---|
| cluster_size | vs_cluster_size |
| cluster_fast | vs_cluster_subseq |
| cluster_unoise | vs_cluster_unoise |
| fastq_join | vs_fastq_join |
| fastq_mergepairs | vs_fastq_mergepairs |
| fastx_subsample | vs_fastx_subsample |
| fastx_filter | vs_fastx_trim_filt |
| fastx_uniques | vs_fastx_uniques |
| search_exact | vs_search_exact |
| sintax | vs_sintax |
| uchime_denovo | vs_uchime_denovo |
| uchime_ref | vs_uchime_ref |
| usearch_global | vs_usearch_global |
| usearch_global (with taxonomic classification (--lcaout)) | vs_alignment_classification |

*Table S2: Number of OTUs at different trimming thresholds before and after chimera removal.*

| **Trimming thresholds** | | **No. of OTUs before chimera removal** | **No. of OTUs after chimera removal** |
|---|---|---|---|
| ***R1*** | ***R2*** | | |
| 230 | 200 | 52 | 45 (86.5%) |
| 236 | 206 | 3 840 | 139 (3.6%) |
| 240 | 210 | 4 065 | 136 (3.3%) |
| 250 | 220 | 3 977 | 121 (3.0%) |
| 260 | 230 | 3 490 | 114 (3.3%) |

#### Supplementary Figures

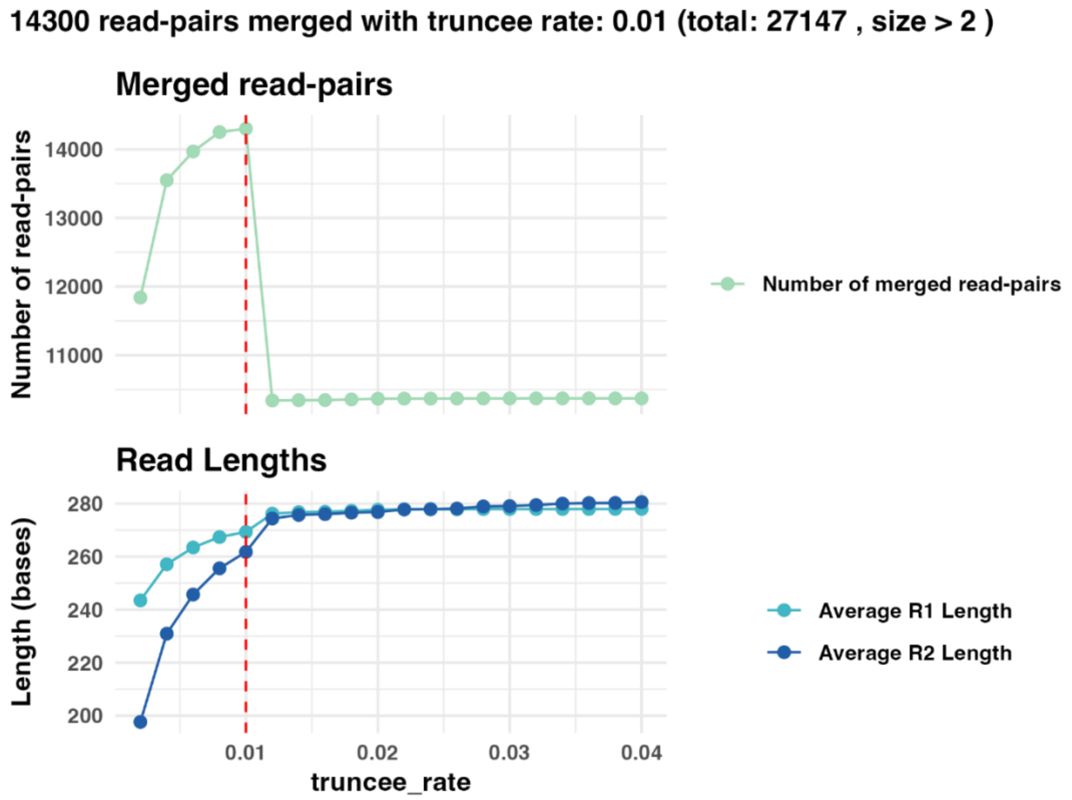


*Figure S1: Optimization of truncee_rate using* `vs_optimize_truncee_rate` *on sample "P2_ZB". The function was run using the default range of* `truncee_rate` *values (0.002 to 0.04). The top panel shows the number of successfully merged read pairs (light green line) with a copy number greater than 2 and a minimum overlap of 10 bases at each setting. The bottom panel displays the average read lengths of R1 (turquoise line) and R2 (dark blue line) reads. The x-axis indicates*

*the tested range of truncee_rate values, and the red vertical line marks the optimal truncee_rate value (0.01). At this setting, 14 300 read pairs were successfully merged.*

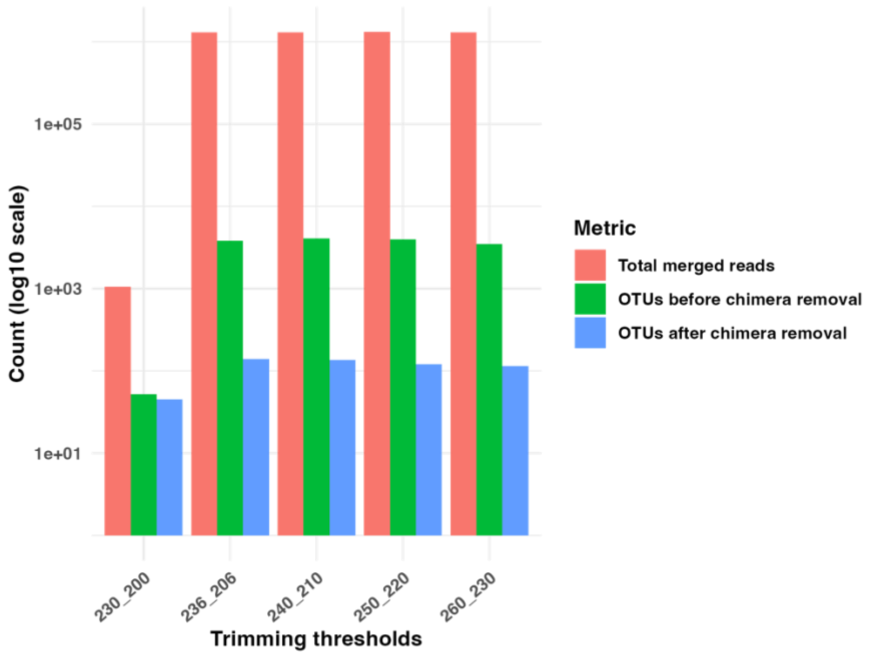


*Figure S2: Comparison of trimming thresholds in DADA2 using the mock community data. Each bar represents different trimming threshold combinations, with the format "R1 length_R2 length". The y-axis (log10 scale) shows the count for the different metrics: total number of merged reads (red), OTUs before chimera removal (green), and OTUs after chimera removal (blue). Trimming thresholds resulting in an overlap length above DADA2's minimum requirement (12 bases) resulted in a comparable number of merged reads. The number of OTUs before chimera removal ranged between approximately 3 400 and 4 100 for these threshold combinations, while the number of OTUs after chimera removal was drastically reduced across all trimming strategies, typically retaining only about 3% of the original OTUs.*
